# Differences in milk microbiota between healthy cows and cows with recurring *Klebsiella* mastitis

**DOI:** 10.1101/2024.02.08.579469

**Authors:** Jingyue Yang, Naiwen Wu, Yindi Xiong, Diego B. Nobrega, Herman W. Barkema, Bingchun Liang, Bo Han, Jian Gao

## Abstract

*Klebsiella* spp. infections continue to have a significant economic impact on the dairy industry, being an important cause of severe clinical mastitis, recurrent infections, and demonstrating poor response to antimicrobials. It is, therefore, essential to investigate the underlying causes of *Klebsiella* spp. infections. Here we used high-throughput DNA sequencing to characterize the milk microbiota of healthy dairy cows (HDC) and cows with history of recurrent *Klebsiella* mastitis (KLB). Our goal was to identify potential pathogenic genera associated with recurrent *Klebsiella* infections in cows. The relative abundance of *Firmicutes* and *Faecalibacterium* was greater in the KLB group than in the HDC group. In contrast, *Proteobacteria* and *Labrenzia* were less abundant than in the HDC group. Although the species distributions differed between groups, diversity and abundance of communities were comparable. Notably, genera of increased occurrence in the KLB group were mostly intestinal-associated, which suggests that cows in the KLB group resided in a contaminated environment or had increased teat-end exposure to fecal bacteria. We did not detect major differences in microbiota among quarters, and also between fore-strip milk and milk collected after fore-stripping. Conversely, milk of heifers had increased alpha diversity in comparison to milk of multiparous cows.

## Introduction

Mastitis is one of the most prevalent diseases in dairy herds, having detrimental impacts on animal welfare and herd economics [1, 2]. Several pathogenic bacteria, including *Klebsiella pneumoniae*, can cause clinical mastitis (**CM**). *Klebsiella* CM can be severe and is often not responsive to antimicrobial therapy [3], and recurrence can occur [4].

When pathogens invade the mammary gland, it is usually followed by a host inflammatory response and changes in the milk composition, which characterizes CM. Importantly, milk from healthy cows also contains diverse bacterial groups, the majority of which are not considered pathogens and unrelated to mastitis. Their role in disease prevention and progression is not fully understood, but recent evidence has demonstrated that the milk microbiota will have an impact in development of mastitis [5]. The advent of high-throughput sequencing technologies has revealed a more significant role of the microbiota in CM than previously thought [6, 7]. The most common method for studying microbiota is to amplify and sequence highly variable regions of 16S rRNA gene copies present in the sample using large-scale sequencing techniques [8].

Analysis of the milk microbiota of dairy cows provides new insights into the ecology of species associated with CM by allowing comparisons between the microbiota of healthy cows and cows with CM [9, 10]. Other applications of large-scale sequencing in mastitis research includes comparisons of the microbiota of clinical and subclinical mastitis [11], changes in milk microbiota during lactation [12] and changes in microbiota before and after antimicrobial treatment [13]. Microbiota analysis can be used to explore mastitis susceptibility and provide new insights as to why some cows develop repeated cases of mastitis and others do not, which can be particularly useful in the context of *Klebsiella* spp. CM.

Here we collected and analyzed milk samples of cows with repeated cases of *Klebsiella* spp. CM and cows with no previous record of CM using 16S rRNA high-throughput sequencing. We performed a comparative analysis of their community structure and diversity to find biomarkers that would be linked to infections.

## Materials and methods

### Farm

This study was conducted in May 2021 on a large commercial dairy farm in the Province of Hebei, China, with 8,832 lactating Holstein cows. Because no human or animal subjects were used, this analysis did not require approval by an Institutional Animal Care and Use Committee or Institutional Review Board. All individuals involved in milk collection were licensed veterinarians with professional veterinary licenses. All procedures were part of routine monitoring and adhered to animal welfare standards. Sampling and publication of this study were conducted with the consent of the dairy farm managers. No animals were sacrificed in the course of this research. The bedding was all recycled fecal solids and animals were fed total mixed rations. Lactating cows were milked three times daily in a circular milking parlor, and the equipment was routinely inspected and sanitized. Commercial iodine-based dipping solutions (0.25% and 0.50% solutions, respectively) were used for pre-and post-milking sanitization. Teats were wiped with a clean towel before cupping. For bacteriological analysis of milk samples, the farm lab makes use of AccuMast Plus® [14].

### Selection of animals and milk sample collection

On-farm clinical records were used to identify cows with history of CM caused by *Klebsiella* spp. Recurring *Klebsiella* spp. cows were defined as having ≥ 2 episodes of *Klebsiella* spp. CM for the current parity, but with no current clinical signs of CM, SCC < 200,000/mL, and no *Klebsiella* spp. growth in milk microbiology culture at the sampling time. Healthy dairy cows (HDC) were lactating cows with no history of CM and SCC <200,000/mL, free of any other clinical disease in the current parity at the time of sampling, and with 4 healthy quarters (e.g., without signs of CM, or anatomically damaged teat endings) [15]. Five cows were selected to represent animals with recurrence of *Klebsiella* spp. mastitis (KLB) and 10 cows with no history of CM were selected as part of the HDC group, including left front (LF), right front (RF), left post (LP), and right post (RP), these four quarters (excluding damaged teats) of each cow, the first three handfuls of milk (FM) and milk collected after forestripping (M) as well as primiparous (P) and multiparous (MP) cows. In each group, 20% of animals were first lactating cows.

Teats were cleaned pre-milking with a 0.50% iodine pre-dip solution, dried with individual paper towels, and then scrubbed for 15 s with a cotton pad soaked in 70% alcohol. The first 3 streaks of milk were collected. Subsequently, the California mastitis test was used in the 4 quarters of each enrolled dairy cow to ensure that all quarters did not have subclinical mastitis. And then milk samples from each quarter of each enrolled cow (approximately 10 - 15 mL) using 50 mL sterile plastic tubes. Samples were put in an incubator with ice packs, preserved at −80°C on the farm, and then transferred to the laboratory for further processing using dry ice.

### DNA extraction and high-throughput sequencing

Total DNA was extracted using the OMEGA Mag-Bind® Soil DNA Kit (M5635-02) following the manufacturer’s instructions. Quality and quantity of DNA were inspected using a NanoDrop ND-2000 spectrophotometer (Thermo Fisher Scientific, Waltham, MA, USA) and agarose gel electrophoresis, respectively. The V3-V4 region of the 16S rRNA gene was amplified by PCR using Q5® High-Fidelity DNA Polymerase with primers 338F (5’-ACTCCTACGGGAGGCAGCA-3’) and 806R (5’-GGACTACHVGGGTWTCTAAT-3’). Agencourt AMPure Beads (Beckman Coulter, Indianapolis, IN) were used to purify PCR amplicons, and the Quant-iT PicoGreen dsDNA Assay Kit was used to measure their quantity (Invitrogen, Carlsbad, CA, USA). Sequencing was performed using the Illumina NovaSeq-PE250 (2 × 250 bp double-ended reads) at Shanghai Personal Biotechnology Co., Ltd (Shanghai, China).

### Bioinformatics analyses

Microbiome bioinformatics was performed using QIIME 2 2019.4 microbiome analysis package [16]. Sequences were quality filtered, denoised, merged, and chimeras removed using the DADA2 plugin to obtain each sample’s amplicon sequence variant (**ASV**) [17]. After acquiring the ASVs, we used the classify-sklearn method of QIIME2 [18] and the nt database (ftp://ftp.ncbi.nih.gov/blast/db/) to annotate species.

Alpha diversity was used to determine the diversity and richness of the microbial community of the samples. The Chao 1 and the Observed species indexes were used to determine species richness, whereas the Shannon and Simpson indexes were utilized to infer species diversity. Beta diversity index focuses on comparing diversity among different habitats, that is, the difference between samples. We utilized the Bray-Curtis distance algorithm to obtain distance matrices among samples, principal component analysis (**PCoA**) and linear discriminant analysis (**LDA**) effect sizes (**LEfSe**) to identify differences between HDC and KLB groups in terms of dominant phylotypes. For testing significance of differences between groups, we used the one-against-all (less strict) comparison strategy to pinpoint species to be compared and Wilcoxon test. Next, the Random Forests algorithm was used to classify samples according to species.

### Statistical analyses

The relative abundance of the major phylum and genus between different strata was determined using GraphPad Prism 8. *P* < 0.05 was considered statistically significant. Comparisons between two groups were done using the Student’s t-test. In contrast, ANOVA were performed to compare results between multiple groups, followed by Tukey’s post hoc test. The error bars in plots indicate the SEM.

## Results

### Descriptive summary

In total, we analyzed 28 milk samples from 5 KLB cows (including M and FM for each sample) and 62 samples from 10 HDC (including M and FM for each sample). The cows’ information is presented in Table 1. A total of 5,184,069 high-quality sequences were retrieved for all samples after using the DADA2 approach for sequence denoising, with a minimum of 37,131 and a maximum of 80,230 sequences. All Illumina sequencing data used in this study can be found under BioProject ID: PRJNA974567.

**Table 1.** Cow information.

| Group | Cow ID | Current parity | Days in milk | The number of clinical mastitis occurrences | Quarters sent for sequencing* |
| --- | --- | --- | --- | --- | --- |
| HDC<sup>1</sup> | 181763 | 1 | 335 | 0 | RF<sup>3</sup> LP RP |
|  | 184557 | 1 | 335 | 0 | LF RF LP |
|  | 177313 | 2 | 271 | 0 | LF RF LP RP |
|  | 156046 | 2 | 271 | 0 | LF RF LP RP |
|  | 141622 | 3 | 155 | 0 | LF RF |
|  | 151354 | 3 | 155 | 0 | LF LP RP |
|  | 158120 | 4 | 381 | 0 | LF RF LP RP |
|  | 141731 | 4 | 364 | 0 | LF RF LP RP |
|  | 140153 | 5 | 228 | 0 | LF RF |
|  | 121480 | 5 | 228 | 0 | LF RF |
| KLB<sup>2</sup> | 139584 | 5 | 338 | 5 | RF LP |
|  | 140607 | 4 | 376 | 6 | LP |
|  | 174030 | 3 | 154 | 7 | LF RF LP RP |
|  | 171695 | 2 | 303 | 7 | LF RF LP RP |
|  | 181699 | 1 | 347 | 2 | LF LP RP |

<sup>1</sup>HDC = healthy dairy cows
<sup>2</sup>KLB = cows with recurrent cases of clinical *Klebsiella* spp. mastitis in this lactation
<sup>3</sup>LF = left front; RF = right front; LP = left post; RP = right post
\* Due to certain cows' broken teats, no four quarters could be collected.

### Milk microbiota vs parity

*Firmicutes* (50.6 - 77.9%), *Proteobacteria* (3.7 - 32.7%), *Bacteroidetes* (8.2 - 14.4%), and *Actinobacteria* (1.5 -17.5%) were identified in primiparous and multiparous cows of both HDC and KLB cows with relative abundance > 1% (Fig 1). Relative abundance of *Firmicutes* (*P* < 0.001) and *Bacteroidetes* (*P* = 0.002) was lower, and that of *Proteobacteria* (*P* < 0.001) was higher in the MP-H group compared to the P-H group. *Actinobacteria* had increased relative abundance in the P-KLB group in comparison to MP-KLB group (*P*< 0.001).

**Fig 1.**
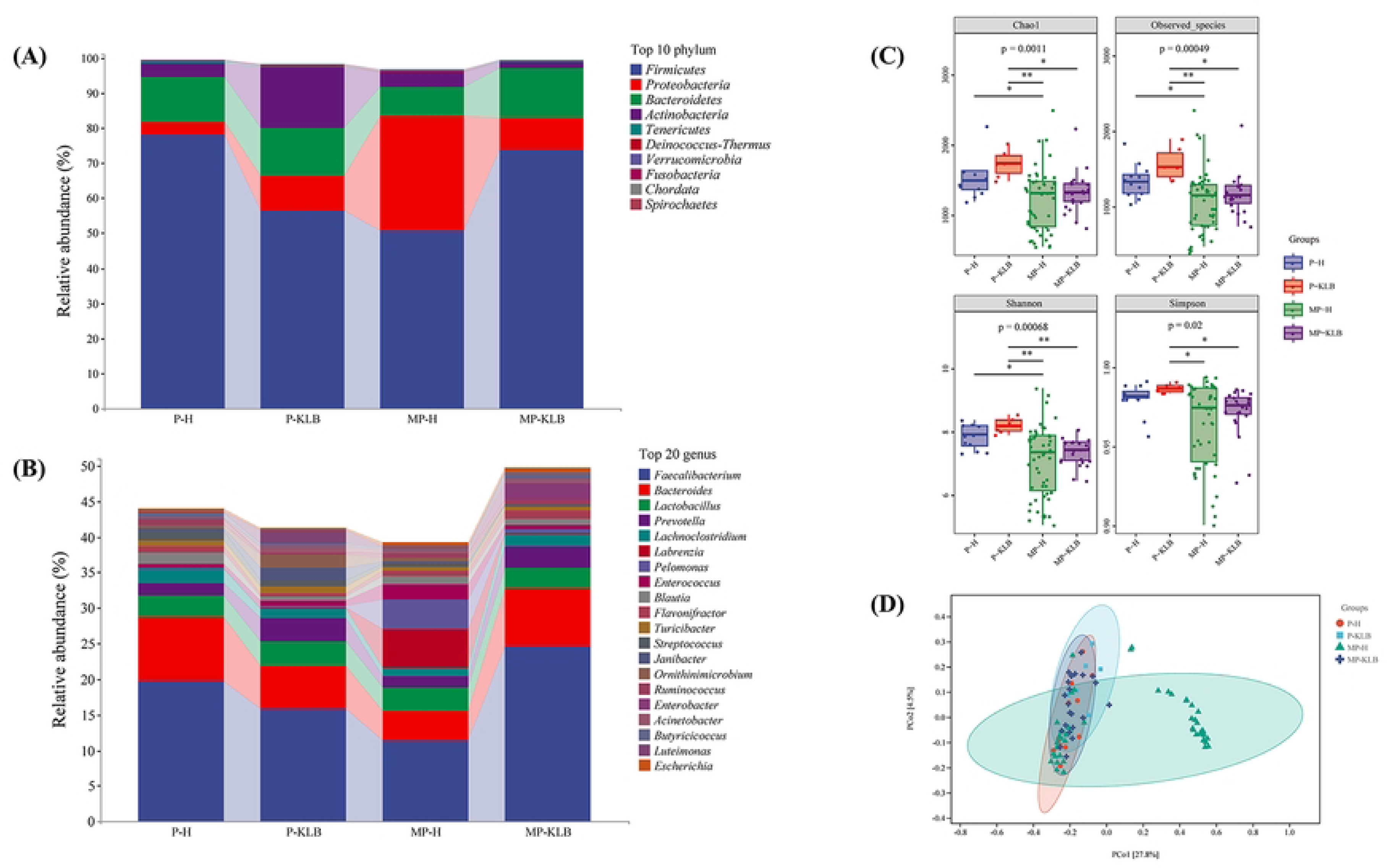
Species diversity and composition at various parities of healthy dairy cows (HDC) and recurrent clinical *Klebsiella* (KLB) mastitis groups. (A) The top 10 phyla in relative abundance, are indicated with each color corresponding to a phylum. (B) Relative abundance of the top 20 genera, with each color corresponding to a different genus. (C) Alpha diversity indices. (D) PCoA analysis of milk microbiota in each group. Coordinates brackets’ percentages represent the proportions of the sample variance data (the distance matrix) that the corresponding coordinate axis can interpret. Each point in the figure represents a sample, and points of different colors indicate different groups. *P-H/KLB = healthy primiparous cow or primiparous cow with recurrent clinical *Klebsiella* spp. Mastitis *MP-H/KLB = healthy multiparous cow or multiparous cow with a recurrent clinical *Klebsiella* spp. mastitis

The P-KLB group had higher relative abundance of *Actinobacteria* (*P*< 0.001) and *Proteobacteria* (*P* = 0.006) whereas the relative abundance of *Firmicutes* (*P*= 0.02) was lower compared to the P-H group. When compared to MP-H, the relative abundance of *Firmicutes* (*P*< 0.001) and *Bacteroidetes* (*P* < 0.001) was higher in the MP-KLB group, while *Proteobacteria* (*P* < 0.001) and *Actinobacteria* (*P*< 0.001) was significantly lower (S1 Table).

The top 20 genera in terms of relative abundance are presented at the genus level (Fig 1B). Only the relative abundance of *Bacteroides* (*P* = 0.001) was lower in group P-KLB than in group P-H, with no other taxa presenting a significant difference between the two groups. *Faecalibacterium* (*P* < 0.001) and *Bacteroides* (*P* < 0.001) had a higher relative abundance in the MP-KLB cows than in MP-H cows, whereas *Labrenzia* (*P*< 0.001) and *Pelomonas* (*P* = 0.004) had a lower relative abundance in MP-KLB cows. In contrast, compared to group P-H, the relative abundance of *Faecalibacterium* (*P* = 0.002) and *Bacteroides* (*P* < 0.001) was lower in group MP-H. The relative abundance of *Faecalibacterium* (*P* = 0.002) and *Bacteroides* (*P* = 0.004) was higher in the P-KLB cows when compared to MP-KLB cows (S2 Table).

The distribution of alpha diversity indices was not significantly different between P-H and P-KLB, and between MP-H and MP-KLB cows (Fig 1C). Conversely, the diversity index pointed that compared with P-KLB, significant lower diversity in milk microbiota in MP-KLB, and the diversity of milk microbiota of healthy primiparous cows was higher than that of healthy multiparous cows.

There was no clear separation of milk samples from the 4 groups within PCoA plots (Fig 1D). The microbial community composition of the 4 groups was relatively similar. Some samples of MP-H clustered separately from others, indicating heterogeneity within the MP-H group.

### Milk microbiota vs quarter location

The relative abundance of genera at the phylum and genus levels did not differ significantly between the 4 quarters of HDC and KLB groups (S3 and S4 Tables). However, we can intuitively see from figs 2A and 2B that at the phylum level, the relative abundance of *Proteobacteria* was higher in the HDC group than in the KLB group for all quarters, and at the genus level, the relative abundance of *Labrenzia* was higher in the HDC group than in the KLB group regardless of quarter location. The different α-diversity indices indicated no significant differences between groups. The PCoA plots showed no clear separation between the groups of samples (Figs 2C and 2D).

**Fig 2.**
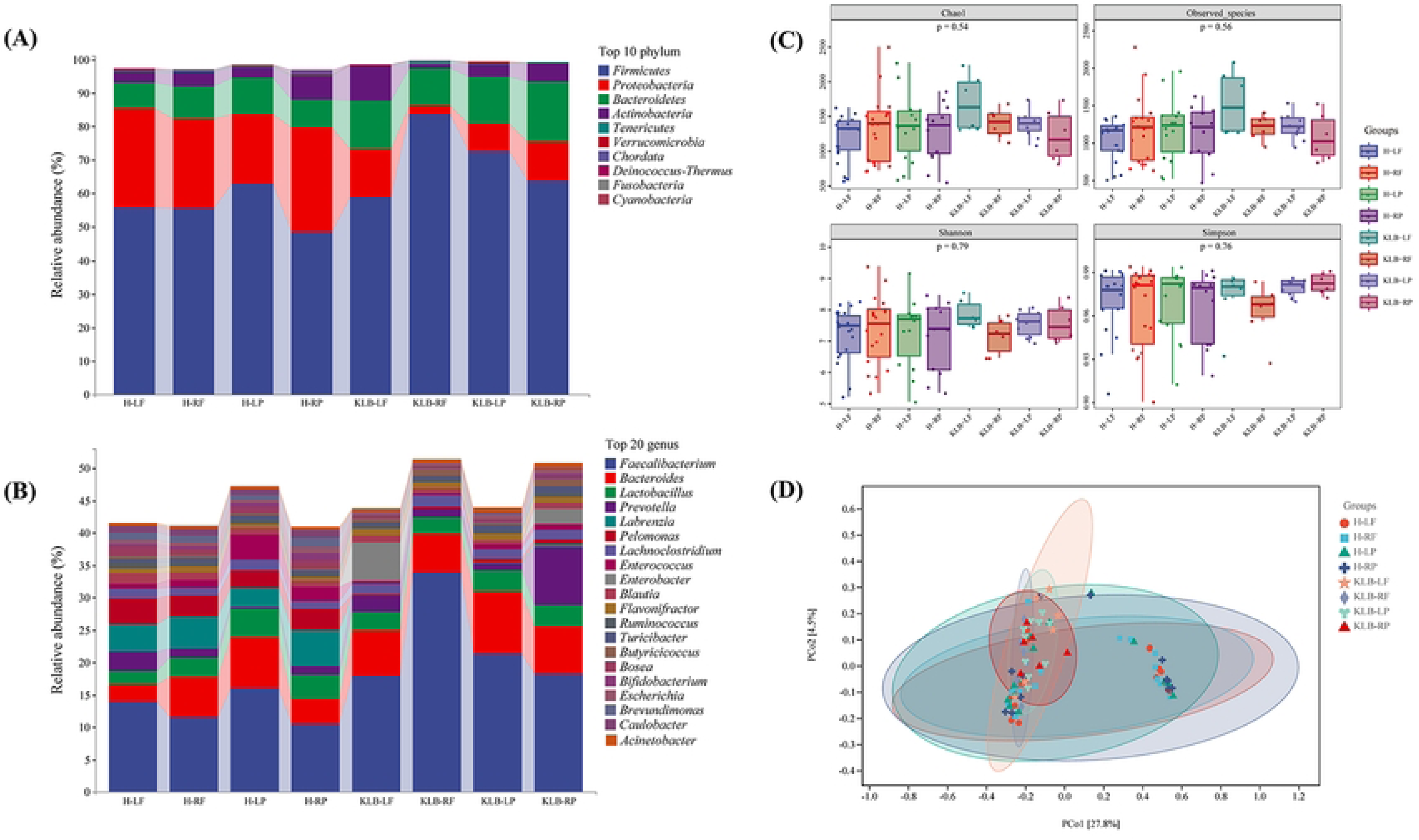
Species diversity and composition at different quarters of healthy dairy cows (HDC) and recurrent clinical *Klebsiella* (KLB) mastitis groups. (A) Top 10 phyla in relative abundance, with each color corresponding to a phylum. (B) Relative abundance of the top 20 genera, with each color corresponding to a different genus. (C) Alpha diversity indices. (D) PCoA analysis of milk microbiota in each group. The coordinates brackets’ percentages represent the proportions of the sample variance data (the distance matrix) that the corresponding coordinate axis can interpret. Each point in the figure represents a sample, and points of different colors indicate different groups. * LF = left front; RF = right front; LH = left hind; RH = right hind

### Milk microbiota: fore-strip milk

We compared the microbiota of the FM and M. The FM in the HDC (FM-H) and KLB (FM-KLB) groups had communities nearly identical to those of the corresponding M groups at the phylum and genus levels (Figs 3A and 3B). For M groups, *Proteobacteria* (*P* = 0.03) were substantially different between M-H and M-KLB at the phylum level (phylum with relative abundance >1%; M-H = 30.8% and M-KLB = 7.3%). For FM groups, *Proteobacteria* had a higher relative abundance in the FM-H group (23.4%) versus the FM-KLB group (11.0%). The relative abundance of the genera in the HDC and KLB groups did not differ significantly at the level of the genus. However, there was a significant difference in the relative abundance of the *Faecalibacterium* (*P* = 0.009), *Pelomonas* (*P*= 0.02) in the M-H group as compared to the M-KLB group (S4 and S5 Tables).

**Fig 3.**
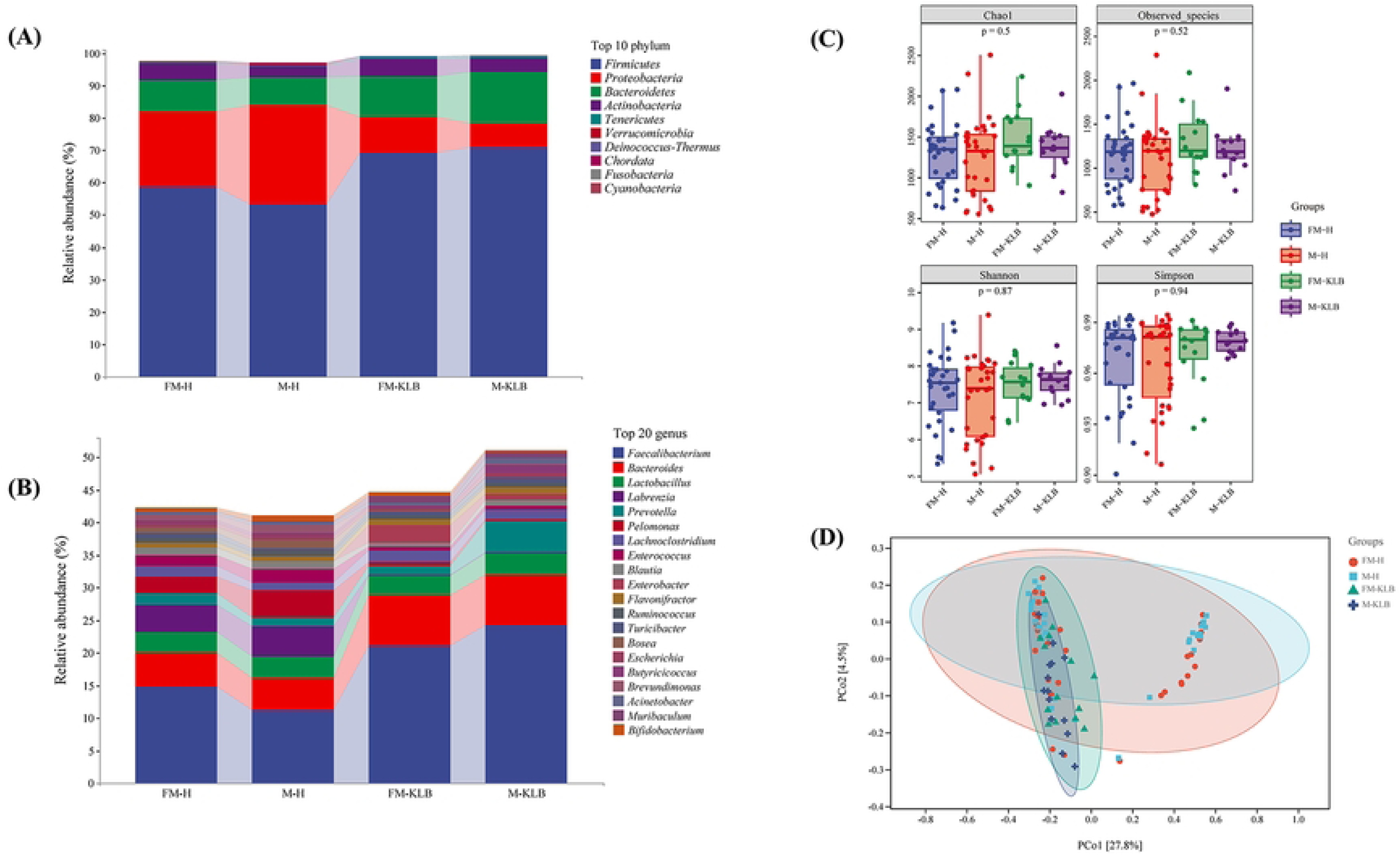
Species diversity and composition between the first three streams of milk and milk collected after forestripping of healthy dairy cows (HDC) and recurrent clinical *Klebsiella* (KLB) groups. (A) Top 10 phyla in relative abundance, with each color corresponding to a phylum. (B) Relative abundance of the top 20 genera, with each color corresponding to a different genus. (C) Alpha diversity indices. (D) PCoA analysis of milk microbiota in each group. The coordinates brackets’ percentages represent the proportions of the sample variance data (the distance matrix) that the corresponding coordinate axis can interpret. Each point in the figure represents a sample, and points of different colors indicate different groups. *FM = first 3 streams of milk; M = milk collected after forestripping

None of the diversity indices significantly differed among groups (Figs 3C and 3D). The PCoA plots demonstrated that all sample areas largely overlapped between the 4 groups.

### Milk microbiota vs *Klebsiella* spp. mastitis

Our previous analyses demonstrated little to no effect of forestripping and quarter location on milk microbiota, whereas diversity indices were influenced by cow parity. Therefore, we used composite milk of multiparous dairy cows for further analysis to control for parity effects. HDC group contained 25 samples, whereas the KLB group contained 11 samples.

Of the top 10 phyla, *Firmicutes* (73.8%) and *Bacteroidetes* (16.8%) were abundant in KLB, whereas *Proteobacteria* (7.0%) and *Actinobacteria* (1.1%) had higher relative abundance in HDC (Figs 4A, 4B and Table 2). The relative abundance of *Faecalibacterium* (26.3%) was the highest in KLB, followed by *Bacteroides* (7.9%) and *Prevotella* (5.6%). In contrast, the relative abundance of *Labrenzia* (0.14%), was lower in the KLB group (*P*= 0.04), and little to no difference between groups was observed for other genus (Figs 5A, 5B and Table 3). Relative abundance of *Faecalibacterium* was substantially higher in KLB than in HDC (*P* < 0.001). **Fig 4**. Top 10 phyla in relative abundance, with (A) each color corresponding to a phylum. (B) Comparison of changes in the proportion of phylum levels with the relative abundance of milk >1% in the healthy dairy cows (HDC) and recurrent clinical *Klebsiella* (KLB) mastitis cows. Error bars indicate the SEM.

**Fig 4.**
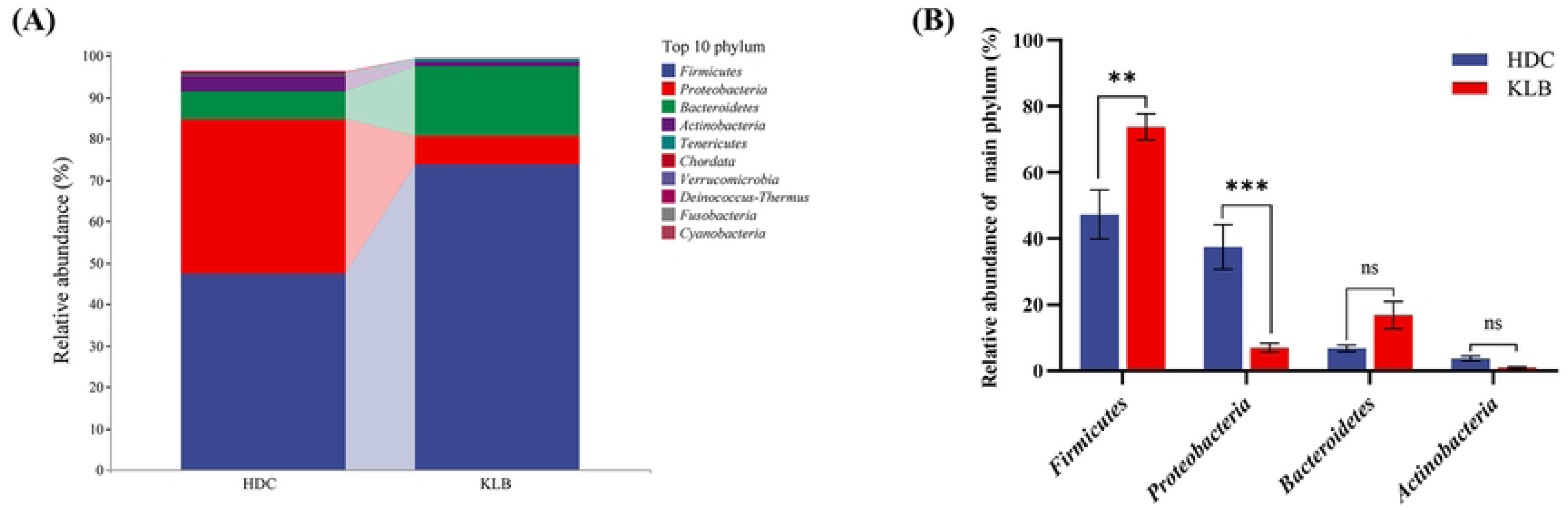
Top 10 phyla in relative abundance, with (A) each color corresponding to a phylum. (B) Comparison of changes in the proportion of phylum levels with the relative abundance of milk >1% in the healthy dairy cows (HDC) and recurrent clinical *Klebsiella* (KLB) mastitis cows. Error bars indicate the SEM.

**Fig 5.**
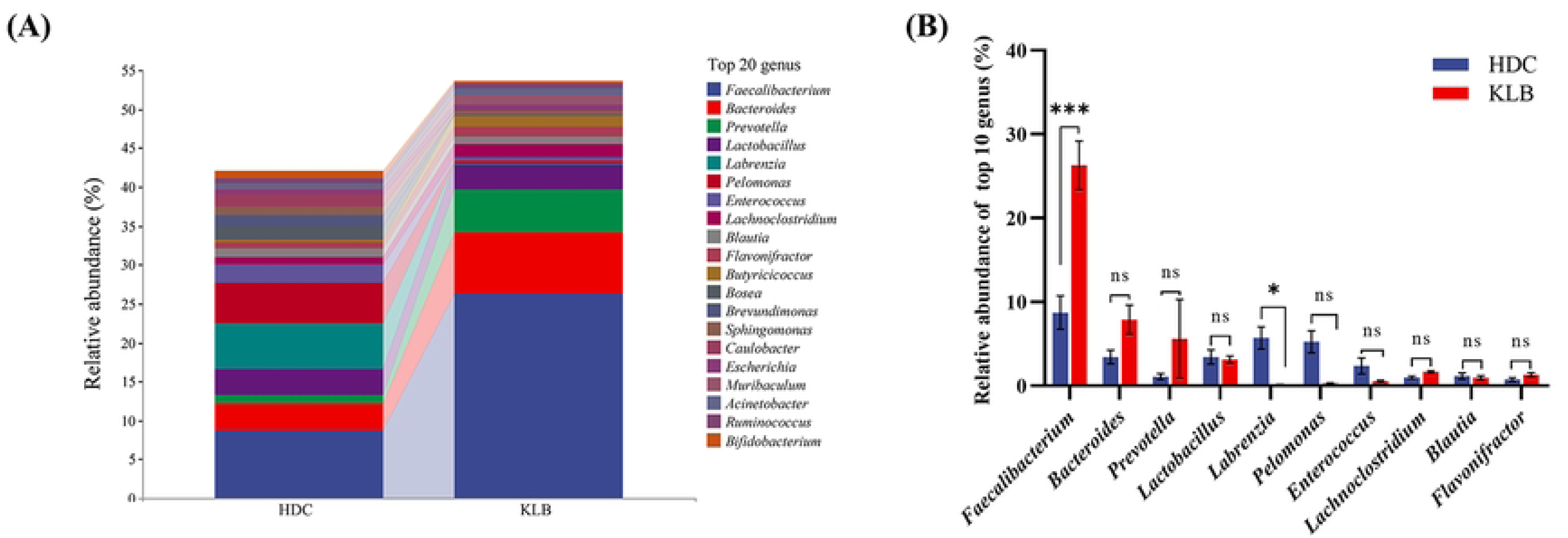
Relative abundance of the top 20 genera, with (A) each color corresponding to a different genus. (B) Comparison of the proportional changes in the levels of the top 10 genera of relative abundance of milk in the healthy dairy cows (HDC) and recurrent clinical *Klebsiella* (KLB) mastitis cows. Error bars indicate the SEM.

**Table 2.** Relative abundance of the top 10 phylum-level microbiota taxa in healthy dairy cows (HDC) and cows with recurrent clinical *Klebsiella* spp. (KLB) mastitis.

| Phylum | Relative abundance (%) |  |
| --- | --- | --- |
|  | HDC | KLB |
| *Firmicutes* | 47.29 | 73.78 |
| *Proteobacteria* | 37.46 | 6.98 |
| *Bacteroidetes* | 6.78 | 16.84 |
| *Actinobacteria* | 3.77 | 1.07 |
| *Tenericutes* | 0.19 | 0.54 |
| *Chordata* | 0.46 | 0.01 |
| *Verrucomicrobia* | 0.2 | 0.16 |
| *Deinococcus-Thermus* | 0.17 | 0.02 |
| *Fusobacteria* | 0.1 | 0.03 |
| *Cyanobacteria* | 0.06 | 0.01 |

**Table 3.** Relative abundance of the top 20 genus-level microbiota taxa in each group.

| Genus | Relative abundance (%) |  |
| --- | --- | --- |
|  | HDC<sup>1</sup> | KLB<sup>2</sup> |
| *Faecalibacterium* | 8.72 | 26.28 |
| *Bacteroides* | 3.43 | 7.88 |
| *Prevotella* | 1.09 | 5.59 |
| *Lactobacillus* | 3.43 | 3.12 |
| *Labrenzia* | 5.71 | 0.14 |
| *Pelomonas* | 5.24 | 0.27 |
| *Enterococcus* | 2.37 | 0.54 |
| *Lachnoclostridium* | 0.99 | 1.65 |
| *Blautia* | 1.15 | 0.96 |
| *Flavonifractor* | 0.75 | 1.29 |
| *Butyricicoccus* | 0.38 | 1.45 |
| *Bosea* | 1.61 | 0.1 |
| *Brevundimonas* | 1.52 | 0.07 |
| *Sphingomonas* | 1.11 | 0.43 |
| *Caulobacter* | 1.46 | 0.05 |
| *Escherichia* | 0.69 | 0.82 |
| *Muribaculum* | 0.15 | 1.25 |
| *Acinetobacter* | 0.6 | 0.75 |
| *Ruminococcus* | 0.69 | 0.63 |
| *Bifidobacterium* | 0.92 | 0.34 |

<sup>1</sup>HDC = healthy dairy cow
<sup>2</sup>KLB = cows with recurrence of clinical *Klebsiella* spp. mastitis

### Diversity indexes vs *Klebsiella* mastitis

Alpha diversity was not significantly different between HDC and KLB (Fig 6A).

**Fig 6.**
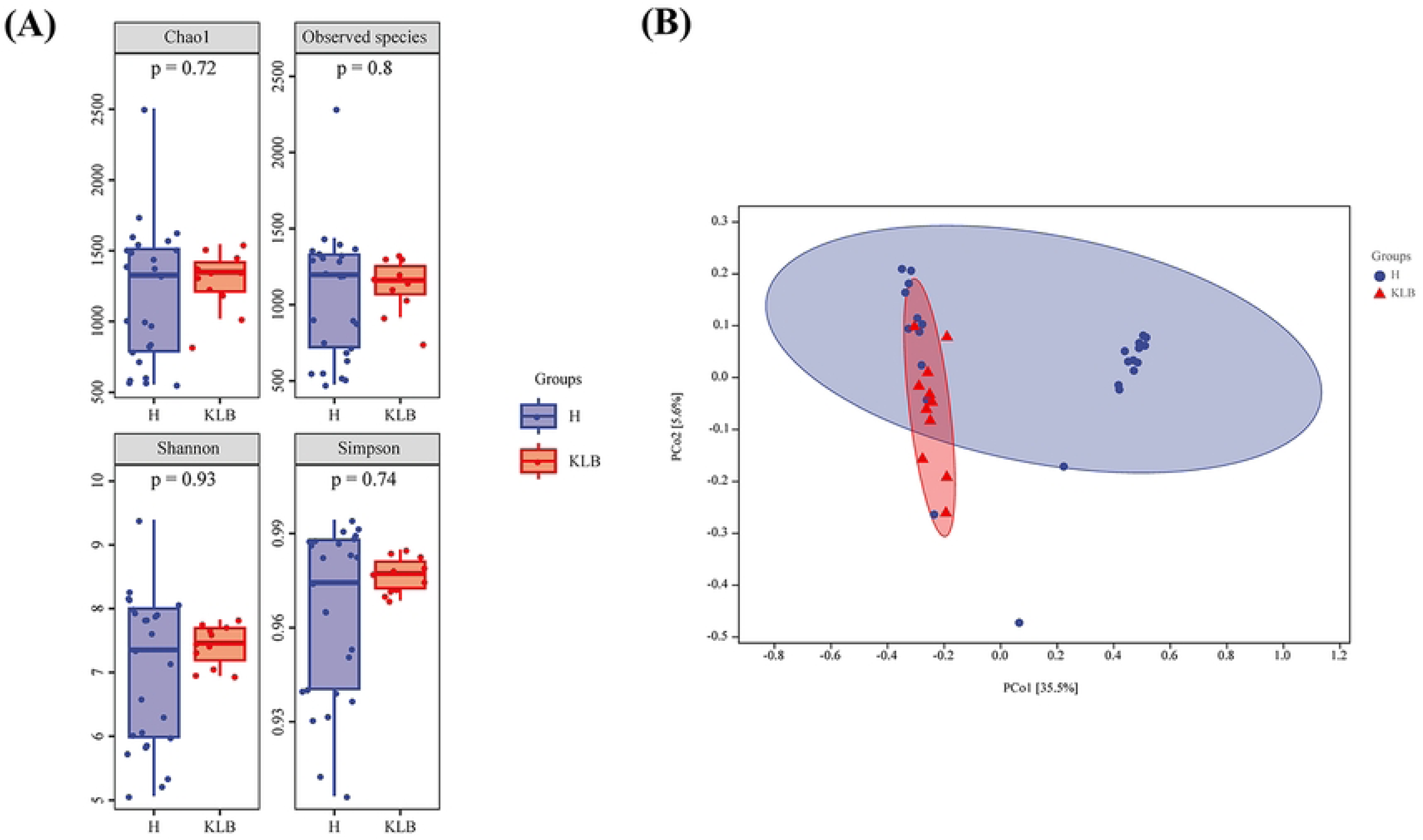
Alpha-diversity indices and PCoA analysis of milk microbiota in each group. (A) The alpha diversity indices, including Chao1, Observed species, Shannon and Simpson, with Chao1 and Observed species indices represent richness indices, and Shannon and Simpson’s indices represent sample diversity. (B) PCoA based on Bray-Curtis. Coordinates brackets’ percentages represent the proportions of the sample variance data (the distance matrix) that the corresponding coordinate axis can interpret. Each point in the figure represents a sample, and points of different colors indicate different groups.

Bray-Curtis distance and PCoA revealed some level of overlap between the 2 groups (Fig 6B), likely explained by individual differences of MP-H cows.

### Compositional differences exist in different groups

HDC and KLB groups had a total of 20,346 ASVs, of which HDC has 12,994 unique ASVs, KLB has 4,741 unique ASVs, and there are 2,611 ASVs in between (Fig 7A). Next, to identify differences in the species composition of the milk microbiome between groups, LEfSe was used to identify biomarkers with a LDA score > 4 (*P* < 0.05). Eight taxa were higher in KLB than HDC, with a high abundance of gut-associated bacteria, such as *Ruminococcaceae*, *Faecalibacterium*, and *Bacteroidetes*. In contrast, the significantly different taxa of HDC compared to KLB is only *Actinobacteria* (Fig 7B). Results of Random Forest sample classifier indicated greater discriminatory power of *Faecalibaculum*, *Faecalibacterium*, *Anaerofustis*, and *Muribaculum* (Fig 7C).

**Fig 7.**
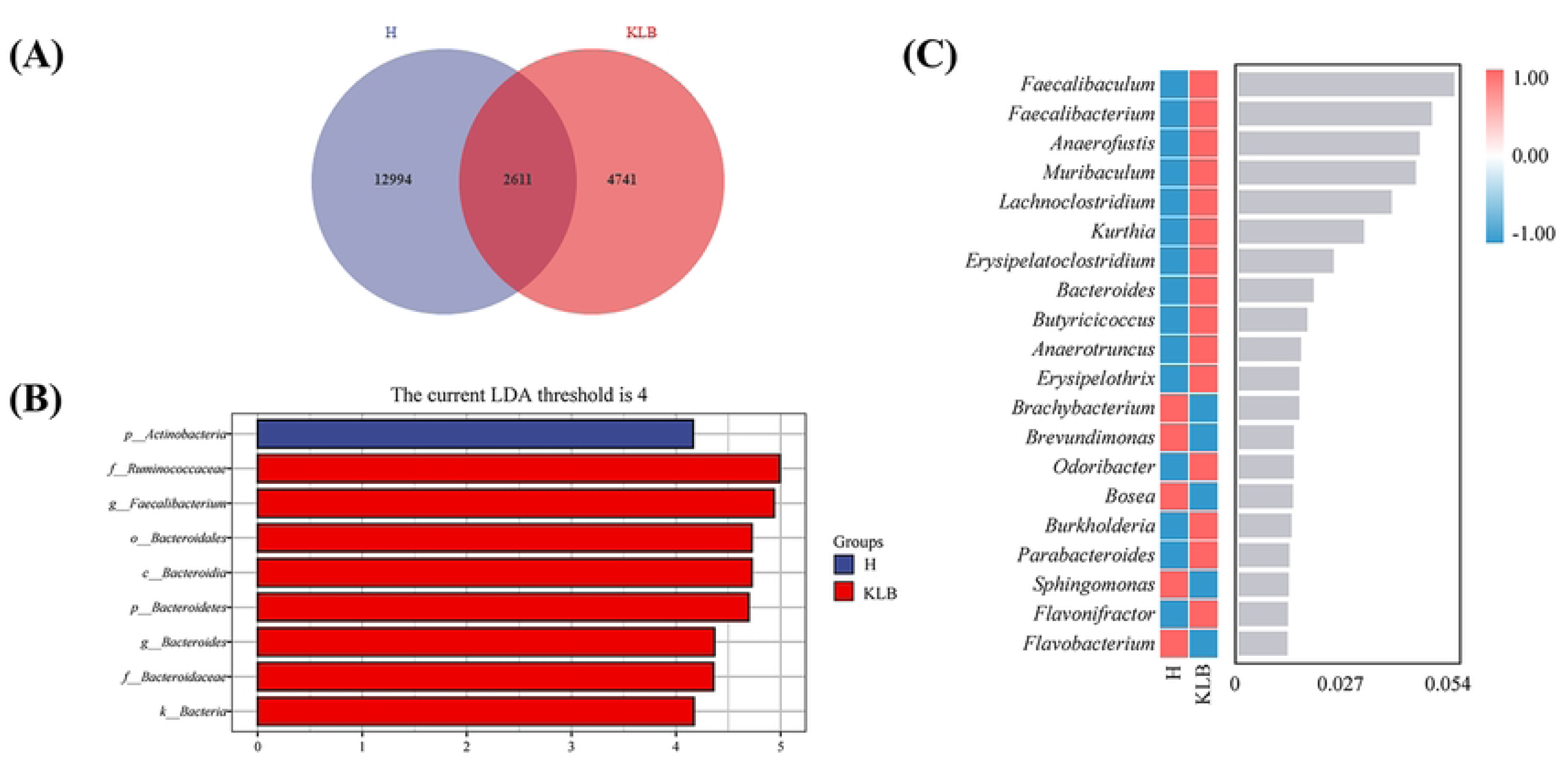
Analysis of species difference and biomarkers. (A) ASVs Venn diagram. Each color block represents a group, the overlapping area between the color blocks indicates the ASVs shared between the corresponding groups, and the number of each block indicates the number of ASVs contained in the block. (B) LEfSe analysis of milk microbiota from 2 groups. The vertical axis presents the taxa with significant differences between groups, whereas the horizontal axis presents the logarithmic score (log 10) of LDA analysis of each taxon in a bar graph. Taxa are sorted by score value to describe their specificity in sample grouping. (C) Random forests analysis of milk microbiota in two groups. The abscissa of the histogram is the scoring value of the importance of the species to the classifier model, and the ordinate is the name of the taxon at the genus level. The heatmap presents the abundance distribution of these species in each group.

## Discussion

Bovine mastitis is one of the major diseases affecting the dairy industry worldwide [19, 20]. Here, we examined the microbial communities of milk samples obtained from the same dairy farm from healthy cows and cows with recurring clinical *Klebsiella* mastitis in the same lactation using high-throughput 16S rRNA gene sequencing. Nonetheless, because some samples’ sequencing quality was subpar and several cows had recurrent *Klebsiella* spp. mastitis that satisfied the screening requirements, resulting in a small sample size for KLB. We also evaluated effects of parity, quarter location, and fore-stripping. Our results contribute to the identification of cow and microbial factors linked with susceptibility to *Klebsiella* spp. CM.

The alpha diversity of the milk community of primiparous cows was higher than that of multiparous cows in both the HDC and KLB cows. Age-dependent changes in microbial compositions affecting the alpha diversity are also seen in the gut of humans [21], and are generally attributed to the diversity of the microbiota of the mother of the newborn fetus. In the analysis of fecal microbiota of primiparous and multiparous cows, it was also found that the fecal microbiota α diversity of primiparous cows was higher than that of multiparous cows [22]. P-H had a larger relative abundance of *Firmicutes* and *Bacteroidetes* and lower abundance of *Proteobacteria* than MP-H, consistent with previous earlier investigations [23].

In the analysis of microbial communities and quarter location, the community diversity was broadly consistent and not different, although there were slight fluctuations at the phylum and genus levels. In a previous study, no differences in microbial composition according to quarter location were detected [23]. As there were no differences in the microbial composition and diversity of milk microbes in the quarters of cows with a history of recurrent *Klebsiella* spp., we can conclude that recurrent *Klebsiella* spp. infections in cows do not affect individual quarter, and that the changes in the microbial structure of milk are uniformly. In one study, the Shannon index was higher in healthy cows than cows with clinical mastitis events, but researchers collected only one quarter per cow [5].

We also compared the microbial community composition between foremilk and milk collected after forestripping, and we found in our results that the microbial communities were similar and consistent in diversity between the first three handfuls of milk and the milk. However, the reason for this situation can not be ruled out that the sample size is too small, so we can subsequently increase the sample size for further testing.

*Firmicutes*, *Proteobacteria*, *Actinobacteria*, and *Bacteroidetes* were the most prevalent bacterial phyla in milk, supporting investigations involving cows, goats and humans [11, 24]. The relative abundance of *Firmicutes* was higher in the KLB group, while the relative abundance of *Proteobacteria* was lower in the KLB group.

*Faecalibacterium* was more abundant in the KLB group than in the HDC group. At the same time, *Labrenzia* was less abundant in the KLB group than in the HDC group. That indicates that *Faecalibacterium* may play an important role in the recovery of diseased cows.

*Faecalibacterium* are commensals in the human gut, and their abundance has been associated with a diet rich in plant-based foods [25, 26, 27]. *Faecalibacterium* were also detected in other studies involving milk microbiomes [28, 29]. *Faecalibacterium prausnitzii* plays a significant role in the immune system, metabolism, inflammation and obesity [30]. In agriculture, *Faecalibacterium prausnitzii* have been tested as probiotics and associated with increased body weight [31, 32]. Our results demonstrated the predominance of *Faecalibacterium* in the KLB group. *Klebsiella* mastitis is an environmental infection that suggests frequent exposure of teats to feces [3]. Increased concentration of *Faecalibacterium* in milk suggests exposure of teats to feces, which is a considerable risk factor for environmental mastitis caused by coliforms.

A new genus of the *Rhodobacteraceae* family, *Labrenzia* (formerly categorized as *Stappia*) has been identified [33]. Roseovarius oyster disease is caused by the bacterial pathogen *Pseudoroseovarius crassostreae* (also known as *Aliiroseovarius crassostreae*). Interestingly, species of this genus such as *Crassostrea virginica*, may protect mollusks from Roseovarius oyster disease [34, 35]. This pathogen negatively impacts both natural oyster populations and oyster-related aquaculture activities. It was proposed that bacterial metabolites may play a role in *Labrenzia spp.*’s protective function. An investigation of the marine bacterium *Labrenzia sp.* 011 reported that this bacterium produces 2 cyclopropane-containing medium-chain fatty acids (1, 2) that inhibit the growth of a range of bacteria and fungi, including the causative agent of Roseovarius oyster disease, *Pseudoroseovarius crassostreae* DSM 16950. In addition, compound 2 acts as a potent partial, β-arrestin-biased agonist on the orphan G-protein-coupled receptor GPR84 activated by medium-chain fatty acids, which is highly expressed in immune cells [36]. However, *Labrenzia* has never been linked to the mammary gland health or milk safety.

Conversely, no significant differences between the two groups regarding the relative abundance of other genera were detected, including for other gut-associated bacteria (*Bacteroides*, *Enterococcus, Prevotella*, *Ruminococcus*, *Blautia*; [37]). Other studies have also demonstrated that *Prevotella*, *Ruminococcus*, and *Bacteroides* can be detected in cow’s milk [28, 38, 39]. These findings indicate that intestinal bacteria are present in milk. Enteric bacteria can reach the mammary gland via the endogenous entero-mammary pathway [40]. However, since the relative abundance of these enteric-associated bacterial genera did not differ among groups, these bacteria may not contribute to the incidence of *Klebsiella* mastitis in dairy cows.

*Acinetobacter* was also detected in both groups. Although their relative abundance was low, studies have demonstrated that *Acinetobacter* is associated with antibiotic resistance and clinical infections in humans [41]. Their occurrence in animal samples may represent a public health concern [41]. Nevertheless, it is unclear if their presence in milk samples represent a risk to farmers and farm personnel.

Diversity and abundance metrics in the HDC and KLB groups were similar, indicating that the microbial diversity and abundance of recovered cows is in line with that of healthy cows. One shortcoming of this study is that milk from cows currently suffering from *Klebsiella* mastitis was not included in the analysis.

We identified biomarkers for the different bacterial taxa in the HDC and KLB groups using lefse analysis. *Actinobacteria* were more abundant in the HDC cows, whereas the enteric-associated genera *Ruminococcaceae*, *Faecalibacterium*, and *Bacteroidales* had higher abundance in the KLB group. This reinforces the importance of the farm environment for controlling *Klebsiella* mastitis. Enteric-associated bacteria were increasingly detected in the milk of KLB cows regardless the presence of clinical disease.

Correspondingly, the random forest analysis indicated that *Faecalibaculum*, *Faecalibacterium*, *Anaerofustis*, *Muribaculum*, and *Lachnoclostridium* can be considered as markers for the KLB cows. *Anaerofustis* also is a genus of gut-associated bacteria. One study investigating changes in the gut microbiota of omnivorous cattle demonstrated a significant decrease in the relative abundance of *Anaerofustis* compared to traditional herbivorous cattle in southern China [42]. In addition, *Anaerofustis stercorihominis*, a species isolated from human feces, plays a role in preventing and treating obesity and metabolic diseases such as diabetes mellitus. However, its role in cows’ health is not well-understood. *Muribaculum* and *Lachnoclostridium* are also gut-associated genera. The increased presence of enteric-associated bacteria in the milk of recovered cows may contribute to recurrent *Klebsiella* infection. This emphasizes that farms should focus on providing a clean environment and bedding to their milking cows to reduce the chances of recurrent mastitis.

Overall, we conclude that milk of cows with history of recurrent clinical *Klebsiella* mastitis had increased presence of intestinal-associated bacteria. Additionally, microbial diversity is associated with parity, and the microbial communities are not different in udder quarters with a different location. Finally, the microbiota of foremilk is similar to that of the remaining milk.

## Supporting information

**S1 Table. Phylum classification of four groups of samples with relative abundance >1%.**

**S2 Table. The top 10 genus levels in relative abundance for the four groups of samples.**

**S3 Table. Phylum classification of 8 groups of samples with relative abundance >1%.**

**S4 Table. The top 10 genus levels in relative abundance for the 8 groups of samples.**

**S5 Table. Phylum classification of four groups of samples with relative abundance >1%.**

**S6 Table. The top 10 genus levels in relative abundance for the four groups of samples.**

The supporting information for this study is all in the “Supporting Information.docx”. All Illumina sequencing data used in this study can be found under BioProject ID: PRJNA974567.

## Author Contributions

**Conceptualization:** Jian Gao.

**Data curation:** Jingyue Yang, Naiwen Wu.

**Formal Analysis:** Jingyue Yang, Yindi Xiong, Yushan Lin.

**Investigation:** Jingyue Yang, Bingchun Liang.

**Supervision:** Jian Gao, Bo Han.

**Visualization:** Jingyue Yang.

**Writing – Original Draft Preparation:** Jingyue Yang.

**Writing – Review & Editing:** Diego B. Nobrega, Herman W. Barkema.

